# Iron depletion has different consequences on the growth and survival of *Toxoplasma gondii* strains

**DOI:** 10.1101/2023.12.21.572787

**Authors:** Eléa A. Renaud, Ambre J.M. Maupin, Yann Bordat, Arnault Graindorge, Laurence Berry, Sébastien Besteiro

**Affiliations:** LPHI, Univ Montpellier, CNRS, 34095 Montpellier, France

**Author notes:** Contact : Sébastien Besteiro. equal contribution (author order was determined randomly) (Eléa A. Renaud, Ambre J.M. Maupin).

**Keywords:** Acute toxoplasmosis, chronic toxoplasmosis, iron depletion, bradyzoites, cystogenic strains

## Abstract

*Toxoplasma gondii* is an obligate intracellular parasite that is responsible for a pathology called toxoplasmosis which is primarily affecting immunocompromised individuals and developing fetuses. The parasite is able to scavenge essential nutrients from its host to support its own growth and survival. Among them, iron is one of the most important elements needed to sustain basic cellular functions, as it is involved in a number of key metabolic processes, including oxygen transport, redox balance and electron transport. We have evaluated the effects of an iron chelator on the development of several parasite strains and found that they differed in their ability to tolerate iron depletion. The growth of parasites usually associated with a model of acute toxoplasmosis was strongly impacted by iron depletion, while cystogenic strains were less sensitive as they were able to convert into persisting developmental forms which are associated with the chronic form of the disease. Ultrastructural and biochemical characterization of the impact of iron depletion on the parasites also highlighted striking changes in both in their metabolism and the one of the host, with a marked accumulation of lipid droplets and perturbation of lipid homeostasis. Overall, our study demonstrates that although acute iron depletion has an important effect on the growth of *T. gondii*, it has a more profound impact on actively dividing parasites, while less metabolically-active parasite forms may be able to avoid some of the most detrimental consequences.

## Introduction

The parasitic protist *Toxoplasma gondii* is responsible for a disease called toxoplasmosis that potentially affects humans and other warm-blooded vertebrates^1^. In immunocompetent individuals, the infection starts with an acute phase, caused by the highly multiplicative and invasive form called tachyzoite, and a long-lasting chronic phase involving an encysted slow-growing form called bradyzoite^2^. In infected immunodeficient individuals, brain-localized bradyzoites may reactivate and this can potentially lead to life-threatening encephalitis. Both developmental stages reside in specific intracellular compartments. Tachyzoites actively invade their host cell and by doing so, they hijack lipids from the host plasma membrane to build a parasitophorous vacuole whose membrane is an important interface supporting parasite survival and replication^3^. Upon initiation of stage conversion into bradyzoites (which can be induced by a number of stresses), the parasitophorous membrane will be heavily modified to build an underlying cyst wall containing various proteins and sugars, providing a protective barrier^4^.

These obligate intracellular parasites have to acquire a number of vital nutrients from their host cells to support their growth and replication^5^, as they have become auxotrophic for important metabolites such as purines or cholesterol, which in fact may be an interesting avenue for the design of novel antiparasitic strategies^6^. One of the key elements that developing parasites need to acquire is iron^7^ which, through its involvement in proteins cofactors like heme or iron-sulfur clusters, is essential to many vital metabolic pathways in the *T. gondii* tachyzoites^8,9^. Maintaining a proper iron balance becomes especially vital in the context of *T. gondii* infection^10,11^ as the parasites need to acquire iron, while the host cells may deprive them of this essential metal. Consequently, host-driven iron withdrawal has the potential to inhibit the growth of these pathogens and can be part of the interferon gamma-dependent response to *T. gondii* infection^12^. During the course of infection, the parasite may also naturally encounter tissues with variable iron availability. For example, the brain typically contains relatively low levels of iron compared to other organs, because excessive iron in the brain can be harmful and is associated with certain neurodegenerative diseases^13^.

Although there is only one *Toxoplasma* species, there is a variety of genotypes that differ in replication rate, the ability to differentiate into bradyzoites, and virulence. The lethality of strains for outbred laboratory mice is the phenotypic marker that has been initially used to define the three archetypal clonal types (I, II and III) of *T. gondii*^14^: type II and III genotypes are usually less virulent and more cystogenic than type I genotypes. These differences in laboratory mouse infections provide a suitable model for mimicking acute (with type I strains) and chronic (with strains of types II and III) infections. In the context of human infections, type II strains are the most predominant among samples in America and Europe, followed by type I and type III (which is only occasionally found in humans)^15^. A wider sampling, both in terms of host and geographical distribution, has subsequently allowed to reveal a much more complex genetic diversity of the parasite^16^. The ability of strains to differentiate into bradyzoites is also a critical point in the efficacy of the treatment as tissue cysts are largely resistant to current drugs impacting tachyzoites.

In this study, we have used the membrane-permeant intracellular Fe^2+^ chelator 2,2′-bipyridine (bipyridyl, BPD) to study the effects of iron deprivation on the survival of *T. gondii* parasites belonging to the I, II and III strains. The chelator had a marked impact on parasite growth and led to specific metabolic changes on both host and parasites, as illustrated by the accumulation of lipid droplets. However, we also noticed that iron depletion led to conversion into the bradyzoite stage for cystogenic parasites belonging to the type II and III strains, which were consequently more likely to survive this metabolic stress.

## Materials and methods

### Parasites and cell culture

Tachyzoites of the RH^17^, Prugniaud (PRU)^18^ and NED^19^ *T. gondii* reference strains were maintained by serial passage in human foreskin fibroblast (HFF, American Type Culture Collection, CRL 1634) cells monolayer or propagated in Vero cells (American Type Culture Collection, CCL 81), grown in Dulbecco’s modified Eagle medium (DMEM, Gibco), supplemented with 5% decomplemented fetal bovine serum, 2-mM L-glutamine and a cocktail of penicillin-streptomycin at 100 μg/ml.

### Plaque assays

Confluent monolayers of HFFs seeded in 24 well plates were infected with freshly egresses tachyzoites. Parasites were seeded in the first lane of the plate and diluted by 1/4 in each subsequent row. The 2,2′-bipyridine (BPD) chelator (D216305, Sigma-Aldrich) was only added 4 hours post-invasion and parasites were subsequently left to grow for 7-10 days for tachyzoites depending on the strain (tachyzoites from type II and II strains have a considerably longer doubling time than those from type I^20^). They were then fixed with 4% v/v PFA and plaques were revealed by staining with a 0.1% crystal violet solution (V5265, Sigma-Aldrich). Pictures of the plaques were acquired with an Olympus MVX10 microscope and plaques area were measured using the ZEN software v2.5 (Zeiss). Plaque areas were expressed relatively to the mean value of those generated by the respective control strain treated with the vehicle (Dimethyl sulfoxide -DMSO-, D5879, Sigma-Aldrich) that was set to 100%.

### IC_50_ determination

IC_50_ values were determined from plaque assays performed with the following concentrations of BPD: 50 µM, 25 µM, 10 µM, 5 µM, 2,5 µM, 1 µM, 0.5 µM. Plaque areas from three independent biological replicates were plotted relative to compound concentrations using the Prism 8 software (Graphpad).

### In vitro conversion to bradyzoites

We used the alkaline pH-induced differentiation^21^ as a control for stage conversion. Briefly, monolayers of HFF grown on coverslips in 24-well plates were infected with 50,000 freshly egressed tachyzoites for 24h in DMEM culture medium. The medium was then replaced by a differentiation medium made of Minimum Essential Medium (MEM) without NaHCO3 (Gibco), supplemented with 50 mM HEPES, 1% penicillin-streptomycin, 1% Glutamine and 3% FBS and adjusted to pH 8.2. Parasites were then cultured at 37°C without CO_2_ and the medium was changed every two days during the whole duration of the experiment.

### Electron microscopy

HFFs infected cells by the different *T. gondii* strains were cultured in Lab-Tek chamber slides (177437, Nunc). Infected cells incubated or not with the chelator were then chemically fixed for 2h at room temperature using the fixation solution (2.5% glutaraldehyde in 0.1 M Cacodylate buffer, pH 7.4). Samples were then stored at 4°C until subsequent processing. Embedding in resin was carried out using a Pelco Biowave PRO+ Microwave processing system (Ted Pella). Program details are provided in Table S1. Samples were post-fixed with 1% osmium tetroxide in 0.1 M cacodylate buffer, pH 7.4. After washing, samples were then incubated in 2% uranyl acetate for 30 minutes at 37°C and further processed in the microwave. After washing, samples were incubated in lead aspartate pre-heated at 50°C. Dehydration was performed with growing concentrations of acetonitrile. Samples were then impregnated in EMbed-812 resin, and polymerized for 48 h at 60°C. All chemicals were from Electron Microscopy Sciences.

Thin serial sections were made using an UCT ultramicrotome (Leica) equipped with an ultra 35° diamond knife (Diatome). Section ribbons were collected on silicon wafers (Ted Pella) for SEM or 100-mesh grids for TEM. Sections on wafers were imaged with a Zeiss Gemini 360 scanning electron microscope on the MRI EM4Bio platform under high vacuum at 1.5 kV. Final images were acquired using the Sense BSD detector (Zeiss) at a working distance between 3.5 and 4 mm. Mosaics were acquired with a pixel size of 5 nm and and a dwell time of 3.2 µs. Sections placed on grids were imaged on a MET LaB6 JEOL 1400 Flash at 100 kV on the Electron Microscopy facility of the university of Montpellier (MEA).

### Immunofluorescence assay

Immunofluorescence assays (IFAs) on tachyzoites and differentiated bradyzoites were performed as described previously^22^. Briefly, monolayers of HFFs grown on coverslips were infected by tachyzoites which were grown for the duration of the experiment with or without addition of BPD, and then fixed with 4% paraformaldehyde in PBS for 20 minutes at room temperature. Cells were washed three times with PBS, permeabilized with 0.3% Triton X-100/PBS for 15 minutes and then saturated with a 1% w/v bovine serum albumin (BSA)/PBS blocking solution for 30 minutes. Proteins were stained with primary antibodies for 1h, followed by three washes with PBS before incubation with secondary antibodies in a 1% BSA/PBS solution for 1h. Primary antibodies used in this study and their respective dilutions were rabbit polyclonal anti-IMC3^23^ antibody diluted at 1/1,000, mouse monoclonal anti-SAG1^24^ diluted at 1/1,000 (T3 1E5), mouse monoclonal anti-P21^25^ diluted at 1/200 (T8 4G10), mouse monoclonal anti-F1-ATPase beta subunit diluted at 1/1,000 (gift of P. Bradley) and rabbit polyclonal anti-pyruvate dehydrogenase E2 subunit^26^ diluted at 1/500. Cyst walls were stained with a 1/300 dilution of a biotin-labelled *Dolichos biflorus* lectin (L-6533, Sigma-Aldrich) for 1h and revealed using a 1/300 dilution of FITC-conjugated streptavidin (SNN1008, Invitrogen). Staining of DNA was performed on fixed cells by incubating them for 5 min in a 1 μg/ml 4,6-diamidino-2-phenylindole (DAPI, 62248, Thermo Fisher) solution. All images were acquired at the MRI facility on a Zeiss Axio Imager Z2 epifluorescence microscope and analyzed with the ZEN v3.6 (Zeiss) and FIJI v1.53t (US National Institutes of Health) softwares. Nile red (72485, Sigma-Aldrich) staining was performed after the fixation and permeabilization steps and prior to antibodies or lectin staining: cells were incubated for 45 minutes with Nile red at a final concentration of 1µg/mL.

### Lipidomic analysis

Parasite extracts for the lipidomic analysis were prepared as follows. Vero cells were grown in 175 cm^2^ flasks until they reached 80% of confluence and were then infected with trachyzoites of the RH strain. 24h after infection, BPD (D216305, Sigma Aldrich) was added at a concentration of 50 µM or not. After 48h of growth for both treated and untreated cell lines, infected host cells were scraped and parasites were released by three passages through a 26G needle. To eliminate cell debris, parasites were filtered through glass wool, then centrifuged washed in ice-cold DPBS (14190-094, Gibco) thrice. After the last wash, 1/10^th^ of the solution was kept to quantify the parasite’s protein content with a bicinchoninic acid assay (UP40840A, Interchim). For each sample, the yield was typically 2.10^8^ parasites corresponding to approximately 800 µg of total proteins. Parasites were finally pelleted down and rapidly frozen before being processed for the lipidomic analyses.

For the quantitative analysis of neutral lipids, lipids were extracted according to Bligh and Dyer^27^ in dichloromethane/methanol/water (2.5:2.5:2, v/v/v), in the presence of the internal standards (stigmasterol, cholesteryl heptadecanoate, glyceryl trinonadecanoate, and DG28). Organic phase was evaporated to dryness and dissolved in 40 µL CH2CL2:MeOH (90:10 v/v). SPE phase was washed with 2X1mL CH2CL2. Neutral lipids were eluted with 2X1 mL CH2CL2:MeOH (90:10,v/v). Samples were dried/recovered in 2X80 µL ethyl acetate. Evaporation was performed and final recovery was in 20 µL ethyl acetate. 1 µL of the lipid extract was analyzed by gas chromatography flam ionization detector on a GC TRACE 1600 Thermo Electron system using an RTX-5 Restek column (5% polysilarylene, 95% polydimethylsiloxane, 5m X 0.25 mm i.d, 0.25 µm film thickness)^28^. Oven temperature was programmed from 190°C to 350°C at a rate of 5°C/min and the carrier gas was hydrogen (5 ml/min). The injector and the detector were at 315°C and 345°C, respectively.

For the quantitative analysis of the phospholipids (PL), samples were extracted according to Bligh and Dyer^27^ in dichloromethane/water/methanol 2% acetic acid (2.5:2:2.5, v/v/v), in the presence 20µL internal standards (TG19) and 40µL internal standards (PC 13:0/13:0 ; Cer d18:1/12:0 ; PE 12:0/12:0 ; SM d18:1/12:0 ; PI 17:0/14:1 ; PS 12:0/12:0) of PL. After centrifugation at 2500 rpm during 6 min, the lipid extract was evaporated to dryness, then dissolved in 50 µL of methanol. PL extracts were analyzed using an Agilent 1290 UPLC system coupled to a G6460 triple quadrupole mass spectrometer (Agilent Technologies) and using MassHunter software for data acquisition and analysis. A Kinetex HILIC column (Phenomenex, 50 x 4.6 mm, 2.6 µm) was used for LC separations. The column temperature was controlled at 40°C. The mobile phase A was Acetonitrile and B was 10 mM ammonium formate in water at pH 3.2. For ceramide, PE, PC, SM, the gradient was as follows: from 10% to 30% B in 10 min; 10-12 min, 100% B; and then back to 10% B at 13 min for 2 min prior to the next injection. The flow rate of mobile phase was 0.3 mL/min and the injection volume was 2 µL. For PI, PS, the gradient was as follows: from 5% to 50% B in 10 min and then back to 5% B at 10.2 min for 9 min prior to the next injection. The flow rate of mobile phase was 0.8 mL/min and the injection volume was 5 µL. Electrospray ionization was performed in positive mode for Cer, PE, PC and SM analysis and in negative mode for PI and PS analysis. Needle voltage was set respectively at 4 kV and −3.5 kV. Analyses were performed in Selected Reaction Monitoring detection mode (SRM) using nitrogen as collision gas. Ion optics and collision energy were optimized for each lipid class. Finally, peak detection, integration and quantitative analysis were done using MassHunter Quantitative analysis software (Agilent Technologies).

### Statistical analyses

Unpaired Student’s t-test was used for comparisons between two groups, and Mann-Whitney non-parametric test was used for particle size comparison (cysts or lipid droplets). They were performed with Prism v8.3 (Graphpad). Unless specified, values are expressed as means ± standard deviation (SD).

## Results

### Iron chelation impacts the growth of types I, II and III of T. gondii

To assess the impact of iron chelation on the growth of *T. gondii* tachyzoites with different cystogenic capacity, we performed plaque assays on type I (RH) and types II (PRU) and III (NED) parasites in the presence of different concentrations of BPD. Plaques are being formed by successive lytic cycles of the parasites (invasion/replication/egress), and thus measuring their area provides a simple and precise assessment of parasite growth. We observed a complete absence of plaques at 50 µM of BPD, and an already important impact on plaque size with 5 µM of BPD (Fig. 1A). The impact at 5 µM was potentially more pronounced for the RH compared with the cystogenic PRU and NED strains, and we thus subsequently performed a detailed assessment of the half maximal inhibitory concentration (IC_50_) of BPD for all strains using increasing concentration ranges of this compounds in plaque assays. We found that the IC_50_ of BPD for the three strains was in the same low micromolar range, although cystogenic strains, with IC_50_ values of about 13 µM, were slightly less impacted than the RH strain, for which the IC_50_ value of BPD was around 7 µM (Fig. 1B).

**Figure 1.**
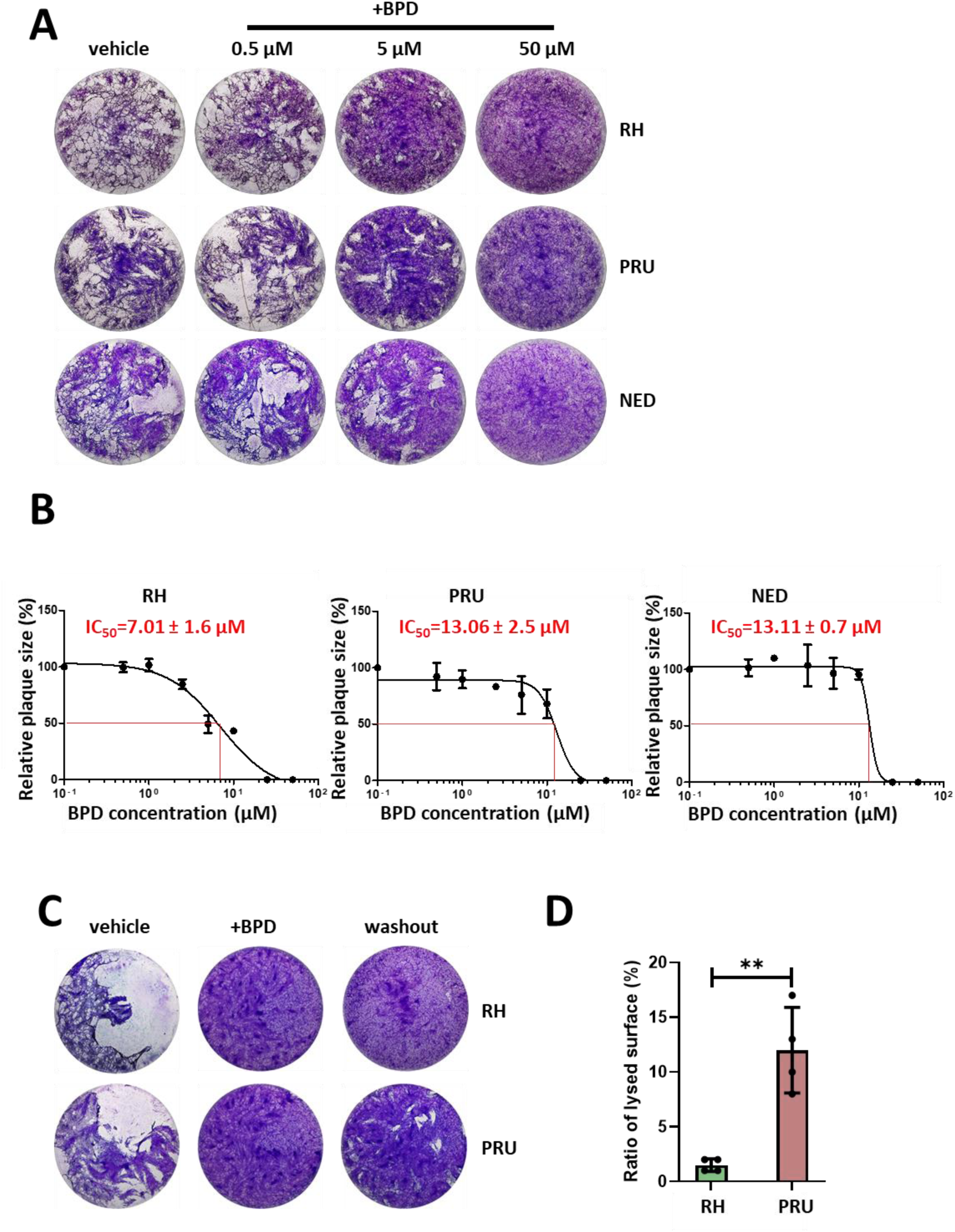
The iron chelator BPD differentially impacts growth of the type I, II and III strains of *T. gondii*. **A.** Plaque assays were performed in the presence or absence of BPD: parasites were added onto HFF monolayer for 7-10 days and lysis plaques were imaged. Included was a control for which the vehicle (dimethyl sulfoxide) only was used. **B.** The half maximal inhibitory concentration (IC_50_) of BPD for type I, II and III strains of *T. gondii* was calculated from plaque assays (as described in A) by assessing relative plaque size compared with the vehicle control. Data are from *n* = 3 independent experiments (except for NED, *n* = 2). Shown are mean values ± SD. **C.** Parasites of the RH and PRU strains were treated for 3 days with 50 µM of BPD, and then treatment continued (+BPD) or the chelator was washed out and parasites were left to grow for another 7 days before imaging of the plaques. **D.** Quantification of the relative total lysed area for the RH and the PRU parasites (compared with the vehicle control) in washout assays performed as described in C. Data are mean values ± SD from *n* = 4 independent experiments. ** denotes *p* ≤ 0.01, Student’s *t*-test.

Plaque assays just allow the global evaluation of parasite fitness, but an absence of plaques does not necessarily reflect death of the parasites, but could also be due to parasites still alive but unable to egress or being largely slowed down in growth. To get further information on the ability of iron depletion to kill tachyzoites of the three strains, we performed a reversibility assay, by incubating the parasites with a high dose of BPD (50 µM) for three days, and then washing out the drug to let them potentially recover for at least a week before measuring plaque size. When performing this on either the RH strain or the cystogenic PRU strain, we noticed that while very few and very little plaques were formed by the former, the latter was able to recover to a significant extent (Fig. 1C, D). This suggests that, although also largely impacted by the BPD treatment, cystogenic strains like the PRU may be able to survive iron starvation more efficiently.

### Acute iron depletion strongly impacts the parasite ultrastructure

We performed immunofluorescence assays (IFAs) on types I, II and III parasites treated for two days with 50 µM BPD to assess the effect of acute iron depletion on the integrity of the parasites. We used markers of the pellicle of the parasite (constituted by the plasma membrane and a double-membrane complex known as the inner membrane complex -IMC-)^29^ and of the mitochondrion and apicoplast, two endosymbiotic organelles that host iron-containing proteins^8,9,26^. We observed important alterations of the pellicle, and problems of DNA or apicoplast segregation in daughter cells (Fig. 2A), however there was no major collapse or fragmentation of the endosymbiotic organelle (Fig. 2A, B). RH parasites seems generally more affected than the parasites of the types II and III strains, especially regarding the integrity of the pellicle outlining the intracellular parasites (Fig. 2A, B).

**Figure 2.**
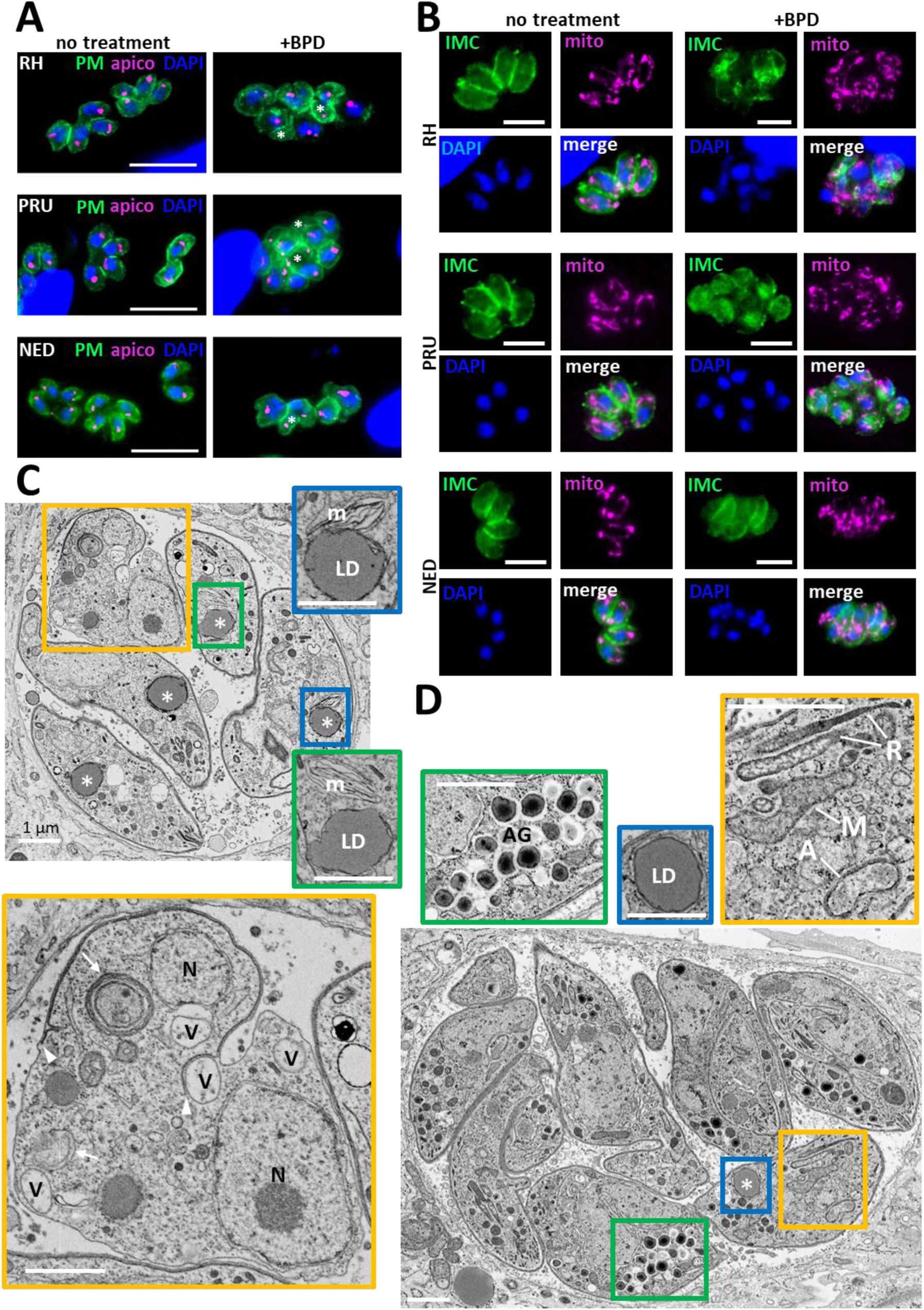
Acute iron chelation by BPD strongly impacts parasite morphology. **A.** Immunofluorescence assay showing co-staining of the parasite plasma membrane (‘PM’, labelled with the anti-SAG1 antibody) and the apicoplast (‘apico’, labelled with the anti-PDH E2 subunit antibody) in parasites from the type I, II and III strains treated or not for 2 days with BPD. Asterisks denote parasites with either nucleus or apicoplast segregation defects. DAPI was used to stain DNA. Scale bar = 10 µm. **B.** Immunofluorescence assay showing co-staining of the parasite inner membrane complex (‘IMC’, labelled with the anti-IMC3 antibody) and the mitochondrion (‘mito’, labelled with the anti-ATPase beta subunit antibody) in parasites from the type I, II and III strains treated or not for 2 days with BPD. DAPI was used to stain DNA. Scale bar = 5 µm. **C.** Electron microscopy analysis of the effects of BPD treatment on the RH strain: asterisks denote lipid droplets (LD) which are magnified on selected insets, along with adjacent multilamellar membranes (m). The main inset shows a dividing parasite with duplicated nuclei (N) that displays interrupted inner membrane complex (arrowheads), as well as many vacuoles (V) and membranous structures potentially resembling autophagic vesicles (arrows). Scale bar = 1 µm. **D.** Electron microscopy analysis of the effects of BPD treatment on the PRU strain: the vacuole contains parasites dividing asynchronously, and displaying structures resembling amylopectin granules (AG, inset), the asterisk denotes a lipid droplet (LD, inset) in a parasite that shows a rather normal aspect for the mitochondrion, rhoptries secretory organelles, apicoplast (M, R and A, in the inset, respectively). Scale bar = 1 µm.

We next used electron microscopy (EM) to assess the impact of BPD treatment on the parasites at the sub-cellular level. Consistent with our IFA observations, we noticed that on parasites of the RH strains displayed organelle segregation problems and a discontinuous IMC (Fig. 2C). In contrast to the synchronized budding and coordinated segregation of organelles that occurs in normal growth conditions, BPD treatment seemed to lead to budding of daughter cells without proper incorporation of nuclear material for instance (Fig. S1). In addition, we could observe vacuolization and accumulation of membranes, which in some instances were surrounding cytosolic material similar to autophagic vesicles^30^. Finally, we observed the frequent presence of large structures resembling lipid droplets in the parasites (LD, Fig. 2C). EM analysis of the effect of BPD on the cystogenic type II PRU strain showed budding problems in some dividing parasites, but there was an overall better preservation of internal membranes and organelles (Fig. 2D). BPD-treated PRU parasites displayed LDs like those found in the RH strain, but in addition we could see structures resembling granules of amylopectin, a storage polysaccharide often found in bradyzoites^31^, suggesting they might be initiating stage conversion (Fig. 2D).

Overall, our data shows that acute treatment with the iron chelator BPD has a strong effect on the growth and morphology of *T. gondii* tachyzoites, but that there might be a different impact on cystogenic or non-cystogenic strains.

### Iron starvation of cystogenic strains leads to an efficient conversion into bradyzoites

As a number of stresses, including nutrient starvation, can initiate the conversion of tachyzoites into bradyzoites and because of our observation that cystogenic strains like the PRU better survive iron depletion (Fig. 1C, D), we next sought to assess more quantitatively the ability of BPD to initiate differentiation. The use of a lectin from the plant *Dolichos biflorus* (DBL), that specifically recognizes a cyst wall glycoprotein called CST1, that is synthesised early upon initiation of stage conversion and accumulates as differentiation progresses^32^. Imaging with fluorescently-labelled DBL allowed identifying the characteristic peripheral labelling of vacuoles having initiated cystogenesis after short term BPD treatment (Fig. 3A). This staining can already be found to occur to some extent in normal *in vitro* culture conditions with cystogenic strains, that usually display some degree of spontaneous differentiation (Fig. 3A, B). However, after two days of BPD treatment, we could see an increased proportion of vacuoles displaying the DBL staining (Fig. 3A, B), although unsurprisingly this was more pronounced in the cystogenic type II and III strains than in the type I strain (with 65, 33 and 22% of DBL-positive vacuoles, respectively).

**Figure 3.**
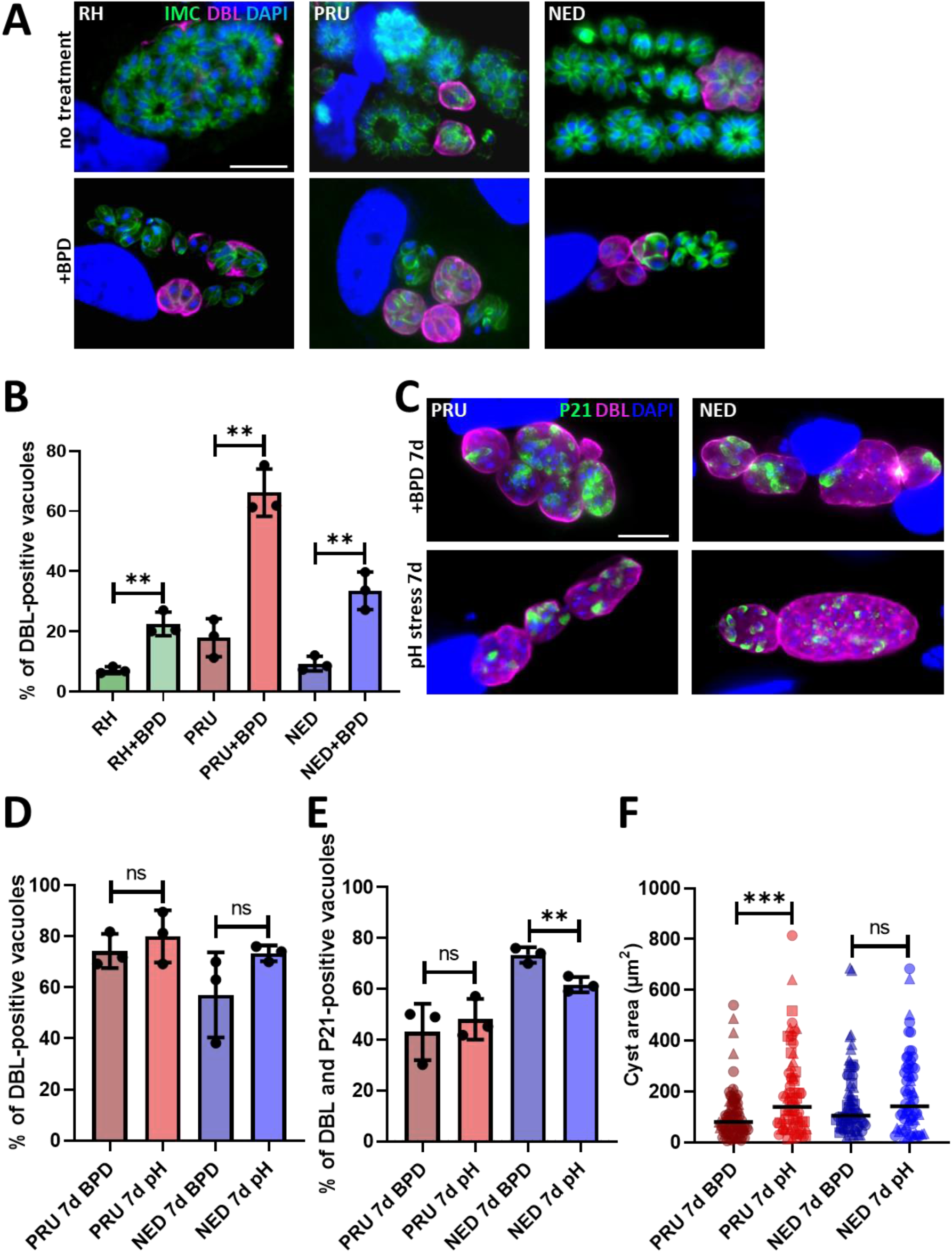
Iron deprivation by BPD triggers stage conversion into bradyzoites. **A.** Parasites from the type I, II and III strains treated or not for 2 days with BPD were stained for the inner membrane complex (IMC) to outline the parasite shape and co-stained with *Dolichos biflorus* lectin (DBL) to detect the maturation of the parasitophorous vacuole membrane into a cyst wall. DAPI was used to stain DNA. Scale bar = 10 µm. **B.** Quantification of DBL-positive vacuoles after 2 days of BPD treatment. Data are mean values ± SD from *n* = 3 independent experiments. At least 50 vacuoles were counted in each experimental condition. ** denotes *p* ≤ 0.01, Student’s *t*-test. **C.** Type II and III cystogenic strains were kept for 7 days in the presence of BPD or stage conversion was induced by alkaline pH stress for the same duration and co-staining was performed for the cysts wall (DBL) and the late bradyzoite marker P21. **D.** Quantification of DBL-positive vacuoles after 7 days of BPD treatment or alkaline pH stress. Data are mean values ± SD from *n* = 3 independent experiments. At least 30 vacuoles were counted in each experimental condition. ns: not statistically significant, Student’s *t*-test. **E.** Quantification of DBL-labelled vacuoles containing P21-positive parasites after 7 days of BPD treatment or alkaline pH stress. Data are mean values ± SD from *n* = 3 independent experiments. At least 20 vacuoles were counted in each experimental condition. ** denotes *p* ≤ 0.01, ns: not statistically significant, Student’s *t*-test. **F.** Measurement of cyst area after 7 days of BPD treatment or alkaline pH stress. Data are mean values ± SD from *n* = 3 independent experiments. At least 25 DBL-positive cysts/vacuoles were measured in each experimental condition. *** denotes *p* ≤ 0.0901, ns: not statistically significant, Student’s *t*-test.

As the BPD washout experiment suggested that cystogenic strains are potentially able to cope better with iron depletion (Fig. 2A, B) and as it may be due to their potential to convert into bradyzoites, we performed long-term incubation with BPD on the PRU and NED strains to assess their ability to convert to more mature cysts. Conversion into bradyzoites is a continuum that happens over the course of several days and while DBL staining is an early marker of cystogenesis, the use of antibodies against P21, a late marker of bradyzoite differentiation whose expression can be detected after 7 days of alkaline pH-induced differentiation^33,34^, is more adequate for the labelling of mature cysts. We performed alkaline pH stress (the most common way to induce conversion to bradyzoites *in vitro*^35^) on the PRU and NED strains in parallel, BPD treatment, both for up to 7 days, and assessed stage conversion by DBL staining and bradyzoite maturation by P21 staining. We observed that both alkaline pH stress and BPD treatment led to comparable conversion rates as assessed by DBL staining (Fig. 3D), and that the proportion of DBL-positive vacuoles that contained P21-positive mature bradyzoites were also similar (Fig. 3C, E). However, the cysts generated upon BPD were slightly smaller than those generated through alkaline pH stress (especially for the PRU strain, Fig. 3F), suggesting that iron chelation has some impact on bradyzoite growth.

Our data thus confirm that iron chelation through BPD treatment leads to an efficient conversion into mature bradyzoites for cystogenic strains of *T. gondii*.

### BPD causes a marked perturbation of lipid homeostasis both in host and parasite cells

When observing by EM the impact at a sub-cellular level of BPD treatment, we noticed an accumulation LD in intracellular parasites (Fig. 2C). The induction of LD was not restricted to the parasites as they were also seen in the host cells and in fact, even in uninfected host fibroblasts, short term (2 days) treatment with BPD led to a marked increase in LD, as seen both by EM and by fluorescence microscopy using Nile Red to stain the LDs (Fig. 4A, B). This is in line with recent findings obtained with several different iron chelators that were shown to induce a quick and robust LD formation in mammalian cells^36–38^. We also observed strong increase in LDs within the parasites of all three strains (Fig. 5A), for which incubation with BPD induced both an increase in LD numbers (Fig. 5B) and size (Fig. 5C).

**Figure 4.**
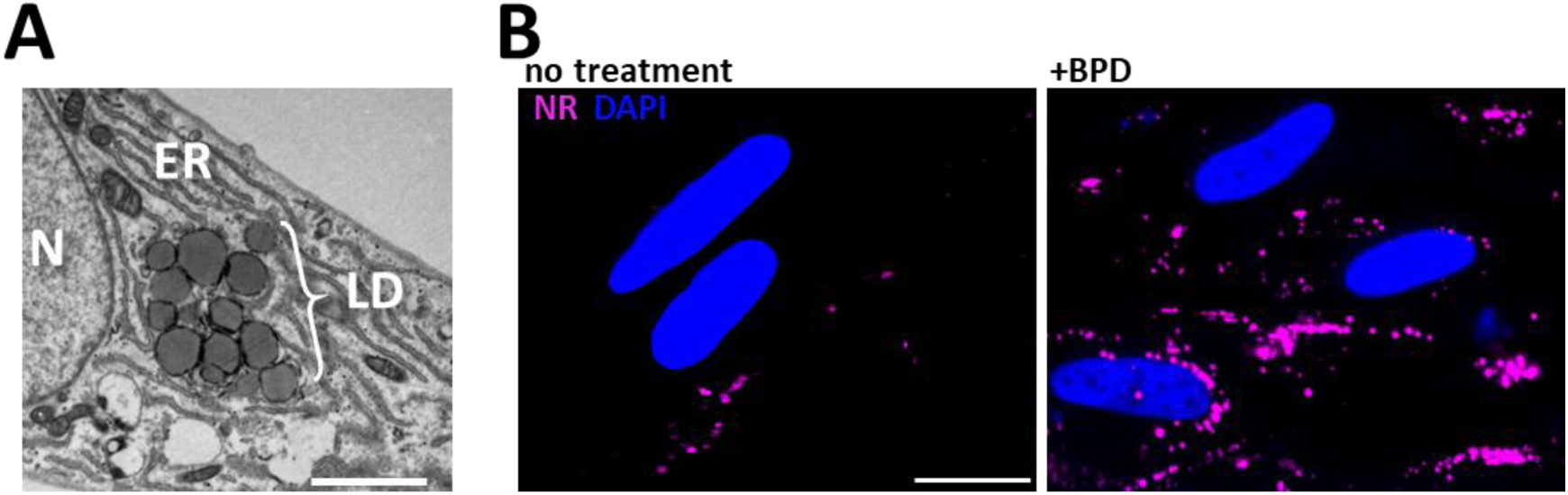
Iron chelation by BPD leads to lipid droplet accumulation in uninfected host cells. **A.** Representative transmission electron microscopy micrograph showing the accumulation of lipid droplets (LD) in a fibroblast after 2 days of BPD treatment. ER: endoplasmic reticulum, N: nucleus. Scale bar = 2 µm. **B.** Fluorescence microscopy picture of fibroblasts either untreated (left) or treated (right) with BPD for 2 days showing that iron chelation leads to an increase in LD, which were labelled with Nile red (NR). DNA was stained with DAPI. Scale bar = 20 µm.

**Figure 5.**
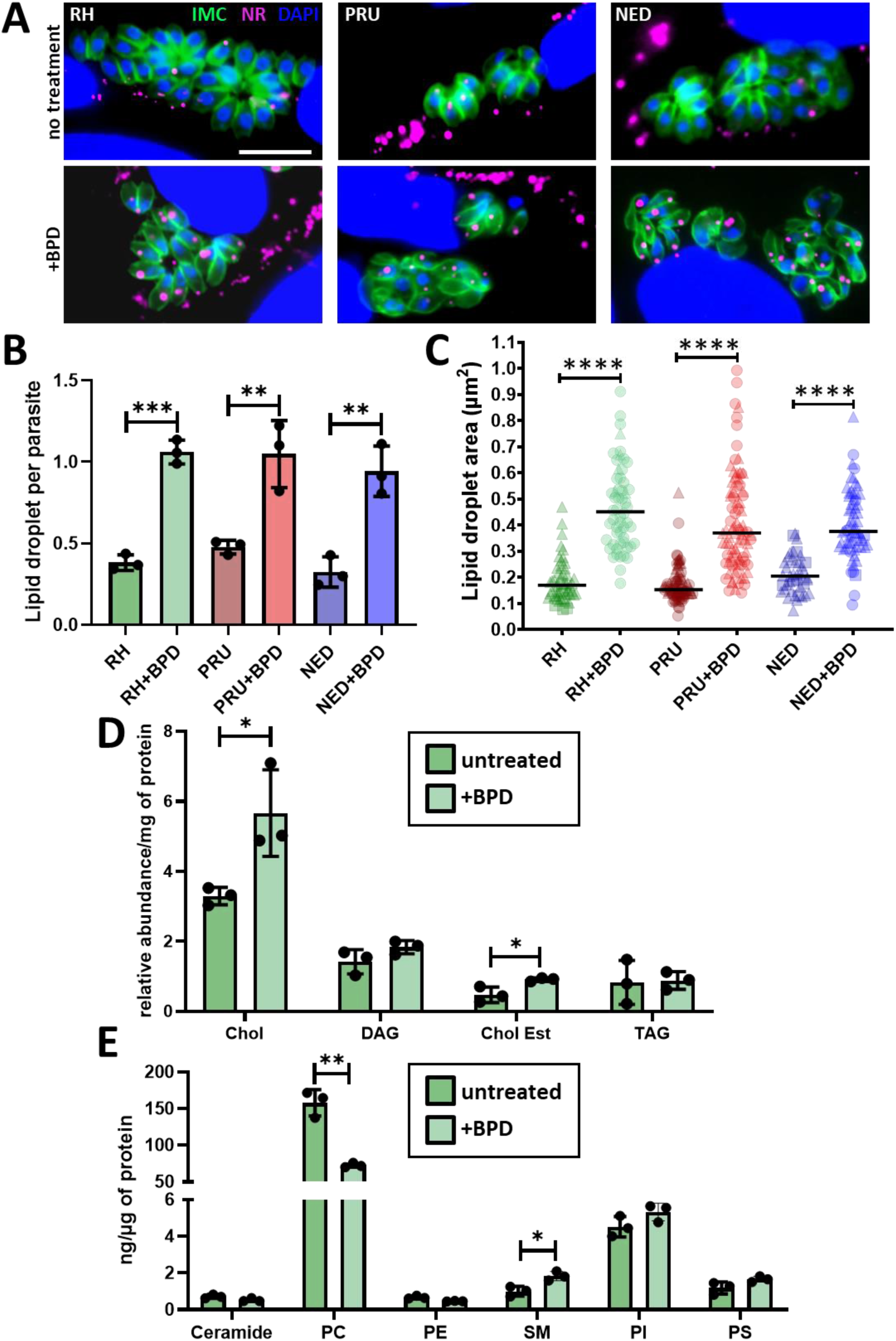
Treatment by BPD affects lipid homeostasis in *T. gondii*. **A.** Intracellular parasites from the type I, II and III strains were treated or not for 2 days with BPD and stained with Nile red (NR) to label the lipid droplets (LD) and counterstained for the inner membrane complex (IMC) protein IMC3 to outline the parasite shape. DNA was stained with DAPI. Scale bar = 10 µm. **B.** Quantification of LD numbers per parasite after treatment or not with BPD for 2 days. Data are mean values ± SD from *n* = 3 independent experiments. At least 280 parasites were counted in each experimental condition. ** denotes *p* ≤ 0.01, *** denotes *p* ≤ 0.001, Student’s *t*-test. **C.** Measurement of LD area in parasites after treatment or not with BPD for 2 days. Data are mean values from *n* = 3 independent experiments. At least 40 LDs were measured in each experimental condition. Symbols are matched between identical experimental groups. **** p ≤ 0.0001, non-parametric Mann-Whitney test. **D.** Analysis of neutral lipid content in parasites after treatment or not with BPD for 2 days. Chol: cholesterol, DAG: diacylglycerol, Chol Est: cholesteryl esters, TAG: triacylglycerol. Data are mean values ± SD from *n* = 3 independent experiments. * denotes *p* ≤ 0.05, Student’s *t*-test. **E.** Analysis of phospholipid content in parasites after treatment or not with BPD for 2 days. PC: phosphatidylcholine, PE: phosphatidylethanolamine, SM: sphingomyelin, PI: phosphatidylinositol, PS: phosphatidylserine. Data are mean values ± SD from *n* = 3 independent experiments. * denotes *p* ≤ 0.05, ** denotes *p* ≤ 0.01, Student’s *t*-test.

The homeostasis of host lipids has important consequences on the one of the parasites, as tachyzoites are able to scavenge lipids stored in host LDs and can incorporate them into its own membranes and LDs^39^. We thus sought to determine the lipidomic profile of RH parasites treated for two days with BPD. LDs are typically composed of a central core of neutral lipids (mostly composed of triacylglycerols and cholesterol esters), surrounded by a monolayer of phospholipids (PLs). The analysis of the parasite’s neutral lipids revealed a marked increase for cholesterol as well as for its fatty acid esters (cholesterol esters), which are typical components of LDs^40^ (Fig. 5D). The analysis of the parasite’s PL content highlighted two significant changes upon BPD treatment: a decrease in phosphatidylcholine and an increase in sphingomyelin (Fig. 5E). Interestingly, phosphatidylcholine is one of the major PLs coating the surface of LDs, and it has been shown previously that a decrease in phosphatidylcholine leads to an increase in LD size^41^ (when the levels of phosphatidylcholine decrease, LDs coalesce via fusion to minimize the surface of the interface and to optimize surface coverage by the PL), so this is perhaps contributing to the enlargement of LDs we seen in addition to their increased numbers after BPD treatment. An increase in sphingomyelin may be attributed to either an increase in scavenging from the host^42^, or an increase in the activity of the parasite’s own sphingomyelin synthase^43^, an enzyme that may both generate sphingomyelin or neutral lipids from phosphatidylcholine, that may then be incorporated into LDs^44^.

The metabolism of long-term persisting bradyzoites relies on the storage of nutrients like carbohydrates^45^, but also potentially lipids^46^. We thus assessed whether or not the marked accumulation of LDs upon iron chelation was due to an indirect effect linked to the differentiation of the parasites. To this end, we assessed LD size and quantification in type II and III bradyzoites obtained after short (3 days) and long (7 days) term differentiation in alkaline pH stress conditions (Fig. S2A). We could see an increase in LD size in converting parasites, especially for type II strain and in particular in late differentiated parasites (Fig. S2C), which is in accordance with a previous study showing that differentiated bradyzoites contain large LDs^46^. However, LDs numbers per parasite did not change (Fig. S2B), which is in sharp contrast with the increase in LDs number induced by BPD treatment (Fig. 5B, Fig. S2A). Moreover, the type I strain is not particularly cystogenic and yet, LDs size and numbers increased in similar proportions as for the cystogenic strains when treated with BPD. Altogether, this suggests that the increase in LDs upon BPD treatment is likely not due to the conversion into bradyzoites, but rather to the more general perturbation of lipid homeostasis caused by iron chelation.

## Discussion

Nutrient limitation, and in particular iron restriction, is a powerful innate immune defense mechanism for host cells to control microbial pathogens^47^. This competition for cellular iron is a delicate balance between both host and microbial systems which is at the heart of the concept of nutritional immunity^48^, with complex implications and consequences that can also be potentially harmful for the host cells themselves. Our results show a very pronounced impact of acute iron depletion on the parasites, with perturbation of membrane homeostasis and defaults in the segregation of organelle and DNA in developing tachyzoites. This is not surprising as the two major classes of cofactors that utilize iron in cells, which are iron-sulfur cluster and heme, are central to proteins that are typically involved in key cellular functions like DNA replication and repair, transcriptional and translational control, respiration, as well as cellular detoxification processes^49^. Not only limiting access to iron seems to be part of the immune response to *T. gondii* infection^12^, but drug-based iron chelation has also been suggested as a potential strategy against the parasite^11^. The impact of iron depletion on *T. gondii* has thus been gaining momentum very recently, with studies investigating for instance in type I parasites the global consequences of iron chelation on the transcriptome of the parasite^50,51^. The main and most obvious trend in the transcriptional profile of iron-depleted *T. gondii* tachyzoites is the induction of the expression of bradyzoite-specific genes^50,51^. Our own quantitative data demonstrates an initiation of stage conversion in all parasite strains we have tested, including type I parasites, but more particularly we have shown that there is a robust induction of differentiation in type II and III cystogenic strains. It was known that reduced availability of nutrients, along with the withdrawal from cell cycle progression or proinflammatory responses for example, can induce tachyzoite-to-bradyzoite differentiation and thus cyst formation^52^. We now show that the use of iron chelators, in spite of having detrimental effect on some parasites, is another tool that can be considered to help generating long term *in vitro* bradyzoites in cystogenic strains, either alone or in jointly with other stress-inducting factors already available^35^.

*T. gondii* is an intracellular parasite that is perfectly adapted to thrive into its host cell, using a combination of *de novo* synthesis and scavenging from the host to obtain essential metabolites^53,54^. Drug-based approaches for interfering with the supply of key elements like iron are affecting both host and parasites and it may be complicated to disentangle the direct from the indirect effects on the parasites. While the recent investigations at the transcriptomic level of the impact of iron chelation on *T. gondii* tachyzoites did not allow to highlight a particular metabolic pathway that would be potentially affected^50,51^, our morphological analysis of BPD-treated parasites showed a pronounced effect on host and parasite lipid homeostasis, as illustrated by the strong accumulation of LDs. *T. gondii* infection results in the recruitment of host LDs and increases biogenesis of these LDs through modification of the host neutral lipid metabolism^55,56^. Recent converging evidence has shown a strong potential for iron chelators to induce LD formation in mammalian cells^36–38^, which we have confirmed in this study. While in mammalian cells it has been suggested that iron depletion induces fatty acid oxidation and leads to mitochondrial dysfunction^38^, the precise molecular mechanisms involved remain essentially unknown. Our lipidomic data suggests that the consequences of iron depletion on the host cell’s lipids subsequently impact the parasite own lipid homeostasis through an increased influx of cholesterol. It is also possible that there is a more direct effect on the parasites of the chelator that would stimulate LD synthesis in similar fashion to the mammalian host cell.

The biogenesis of LDs is part of an integrated stress response to cellular injuries, and not only their biogenesis is quite logically induced in cells exposed to excess amounts of lipids, but also in conditions of acute nutrient deprivation and after oxidative stress^57^. LDs contribute to the protection of cellular integrity by buffering the excess of potentially toxic lipids, maintaining energy and redox balance and preserving membrane homeostasis. As confirmed by our microscopic observation, acute iron depletion led to a pronounced impact on the parasite’s membrane homeostasis, and the strong increase in LD formation in the parasite may not only be a consequence of the increased amount of lipids in the host, but also as a part of the parasite’s own stress response mechanism. *T. gondii* tachyzoites mitigate stress damage through an integrative stress response pathway that can for example be activated in conditions of nutrient starvation^58^ or acute oxidative stress^59^, and leads to stage conversion into the persistent bradyzoite form^60^. Conversion from fast-replicating tachyzoites to this cyst-confined developmental form is induced by a variety of stresses and, thanks to a very slow replication and a reduced metabolic activity, extends survival in adverse conditions^35^. Our results demonstrate that while acute iron depletion strongly and rapidly impacts the viability of type I parasites, it leads to a relatively efficient conversion into the bradyzoite form for the type II and III cystogenic strains. Long term acute iron deprivation may nevertheless also be detrimental for bradyzoites, but such a treatment will likely also be damaging for the host cells. In the mammalian host, bradyzoites persist in specific tissues that include the brain where there is an overall low amount of this element^13^ (although there are probably regional iron concentration differences^61^). It is thus possible that these forms are more adapted to respond to low iron availability. Bradyzoites are largely resistant to the current therapies used in the context of acute toxoplasmosis. Our results show the ability of cystogenic strains to efficiently convert into bradyzoites upon treatment with an iron chelator, and to persist in these conditions for several days *in vitro*. Thus, concerning the perspective of using iron deprivation as a treatment for treating toxoplasmosis^11^, our findings are a cautionary note that it might not be particularly efficient in the context of chronic toxoplasmosis.

## Acknowledgements

Thanks to J.F. Dubremetz, M.L. Dardé, P. Bradley and M.J. Gubbels for strains and antibodies. We thank the “Montpellier ressources imagerie” (MRI) platform and the Electron Microscopy facility of the university of Montpellier for providing access to their microscopes. Lipidomic analyses were performed on the “MetaToul-Lipidomique” platform (I2MC, Inserm, Toulouse, France - MetaboHUB-ANR-11-INBS-0010). This work was supported by the Agence Nationale de la Recherche (grant ANR-22-CE20-0026).

## References

1. Montoya JG, Liesenfeld O. Toxoplasmosis. Lancet 2004; 363:1965–76.

2. Sanchez SG, Besteiro S. The pathogenicity and virulence of Toxoplasma gondii. Virulence 2021; 12:3095–114.

3. Clough B, Frickel E-M. The Toxoplasma parasitophorous vacuole: an evolving host– parasite frontier. Trends in Parasitology 2017; 33:473–88.

4. Tu V, Tomita T, Sugi T, Mayoral J, Han B, Yakubu RR, Williams T, Horta A, Ma Y, Weiss LM. The Toxoplasma gondii cyst wall interactome. mBio 2020; 11:e02699–19,/mbio/11/1/mBio.02699-19.atom.

5. Saliba KJ, Kirk K. Nutrient acquisition by intracellular apicomplexan parasites: staying in for dinner. International Journal for Parasitology 2001; 31:1321–30.

6. Coppens I. Exploitation of auxotrophies and metabolic defects in Toxoplasma as therapeutic approaches. International Journal for Parasitology 2014; 44:109–20.

7. Sloan MA, Aghabi D, Harding CR. Orchestrating a heist: uptake and storage of metals by apicomplexan parasites. Microbiology (Reading) 2021; 167:mic.0.001114.

8. Bergmann A, Floyd K, Key M, Dameron C, Rees KC, Thornton LB, Whitehead DC, Hamza I, Dou Z. Toxoplasma gondii requires its plant-like heme biosynthesis pathway for infection. PLoS Pathog 2020; 16:e1008499.

9. Pamukcu S, Cerutti A, Bordat Y, Hem S, Rofidal V, Besteiro S. Differential contribution of two organelles of endosymbiotic origin to iron-sulfur cluster synthesis and overall fitness in Toxoplasma. PLoS Pathog 2021; 17:e1010096.

10. Aghabi D, Sloan M, Gill G, Hartmann E, Antipova O, Dou Z, Guerra AJ, Carruthers VB, Harding CR. The vacuolar iron transporter mediates iron detoxification in Toxoplasma gondii. Nat Commun 2023; 14:3659.

11. Oliveira MC, Coutinho LB, Almeida MPO, Briceño MP, Araujo ECB, Silva NM. The availability of iron is involved in the murine experimental Toxoplasma gondii infection outcome. Microorganisms 2020; 8:560.

12. Dimier IH, Bout DT. Interferon-gamma-activated primary enterocytes inhibit Toxoplasma gondii replication: a role for intracellular iron. Immunology 1998; 94:488–95.

13. Ward RJ, Zucca FA, Duyn JH, Crichton RR, Zecca L. The role of iron in brain ageing and neurodegenerative disorders. The Lancet Neurology 2014; 13:1045–60.

14. Howe DK, Sibley LD. Toxoplasma gondii comprises three clonal lineages: correlation of parasite genotype with human disease. J Infect Dis 1995; 172:1561–6.

15. Hosseini SA, Amouei A, Sharif M, Sarvi S, Galal L, Javidnia J, Pagheh AS, Gholami S, Mizani A, Daryani A. Human toxoplasmosis: a systematic review for genetic diversity of Toxoplasma gondii in clinical samples. Epidemiol Infect 2018; 147:e36.

16. Galal L, Hamidović A, Dardé ML, Mercier M. Diversity of Toxoplasma gondii strains at the global level and its determinants. Food and Waterborne Parasitology 2019; 15:e00052.

17. Sabin AB. Toxoplasmic encephalitis in children. JAMA 1941; 116:801.

18. Martrou P, Pestre, M, Loubet R, Nicolas, Malinvaud G. La toxoplasmose congénitale (note concernant un cas mortel). Limousin Médical 53:3–7.

19. Cristina N, Dardé ML, Boudin C, Tavernier G, Pestre-Alexandre M, Ambroise-Thomas P. A DNA fingerprinting method for individual characterization ofToxoplasma gondii strains: combination with isoenzymatic characters for determination of linkage groups. Parasitol Res 1995; 81:32–7.

20. Radke J. Defining the cell cycle for the tachyzoite stage of Toxoplasma gondii. Molecular and Biochemical Parasitology 2001; 115:165–75.

21. Soete M, Camus D, Dubremetz JF. Experimental induction of bradyzoite-specific antigen expression and cyst formation by the RH strain of Toxoplasma gondii in vitro. Experimental Parasitology 1994; 78:361–70.

22. Sanchez SG, Bassot E, Cerutti A, Mai Nguyen H, Aïda A, Blanchard N, Besteiro S. The apicoplast is important for the viability and persistence of Toxoplasma gondii bradyzoites. Proc Natl Acad Sci U S A 2023; 120:e2309043120.

23. Anderson-White BR, Ivey FD, Cheng K, Szatanek T, Lorestani A, Beckers CJ, Ferguson DJP, Sahoo N, Gubbels M-J. A family of intermediate filament-like proteins is sequentially assembled into the cytoskeleton of Toxoplasma gondii. Cell Microbiol 2011; 13:18–31.

24. Couvreur G, Sadak A, Fortier B, Dubremetz JF. Surface antigens of Toxoplasma gondii. Parasitology 1988; 97 (Pt 1):1–10.

25. Tomavo S, Fortier B, Soete M, Ansel C, Camus D, Dubremetz JF. Characterization of bradyzoite-specific antigens of Toxoplasma gondii. Infect Immun 1991; 59:3750–3.

26. Renaud EA, Pamukcu S, Cerutti A, Berry L, Lemaire-Vieille C, Yamaryo-Botté Y, Botté CY, Besteiro S. Disrupting the plastidic iron-sulfur cluster biogenesis pathway in Toxoplasma gondii has pleiotropic effects irreversibly impacting parasite viability. J Biol Chem 2022; 298:102243.

27. Bligh EG, Dyer WJ. A rapid method of total lipid extraction and purification. Can J Biochem Physiol 1959; 37:911–7.

28. Barrans A, Collet X, Barbaras R, Jaspard B, Manent J, Vieu C, Chap H, Perret B. Hepatic lipase induces the formation of pre-beta 1 high density lipoprotein (HDL) from triacylglycerol-rich HDL2. A study comparing liver perfusion to in vitro incubation with lipases. J Biol Chem 1994; 269:11572–7.

29. Morrissette NS, Murray JM, Roos DS. Subpellicular microtubules associate with an intramembranous particle lattice in the protozoan parasite Toxoplasma gondii. J Cell Sci 1997; 110 (Pt 1):35–42.

30. Besteiro S, Brooks CF, Striepen B, Dubremetz J-F. Autophagy protein Atg3 is essential for maintaining mitochondrial integrity and for normal intracellular development of Toxoplasma gondii tachyzoites. PLoS Pathog 2011; 7:e1002416.

31. Dubey JP, Lindsay DS, Speer CA. Structures of Toxoplasma gondii tachyzoites, bradyzoites, and sporozoites and biology and development of tissue cysts. Clin Microbiol Rev 1998; 11:267–99.

32. Tomita T, Bzik DJ, Ma YF, Fox BA, Markillie LM, Taylor RC, Kim K, Weiss LM. The Toxoplasma gondii cyst wall protein CST1 is critical for cyst wall integrity and promotes bradyzoite persistence. PLoS Pathog 2013; 9:e1003823.

33. Dogga SK, Lunghi M, Maco B, Li J, Claudi B, Marq J-B, Chicherova N, Kockmann T, Bumann D, Hehl AB, et al. Importance of aspartyl protease 5 in the establishment of the intracellular niche during acute and chronic infection of Toxoplasma gondii. Mol Microbiol 2022;

34. Soete M, Fortier B, Camus D, Dubremetz JF. Toxoplasma gondii: kinetics of bradyzoite-tachyzoite interconversion in vitro. Exp Parasitol 1993; 76:259–64.

35. Cerutti A, Blanchard N, Besteiro S. The bradyzoite: a key developmental stage for the persistence and pathogenesis of toxoplasmosis. Pathogens 2020; 9.

36. Pereira M, Chen T-D, Buang N, Olona A, Ko J-H, Prendecki M, Costa ASH, Nikitopoulou E, Tronci L, Pusey CD, et al. Acute iron deprivation reprograms human macrophage metabolism and reduces inflammation in vivo. Cell Rep 2019; 28:498–511.e5.

37. Crooks DR, Maio N, Lane AN, Jarnik M, Higashi RM, Haller RG, Yang Y, Fan TW-M, Linehan WM, Rouault TA. Acute loss of iron-sulfur clusters results in metabolic reprogramming and generation of lipid droplets in mammalian cells. J Biol Chem 2018; 293:8297–311.

38. Long M, Sanchez-Martinez A, Longo M, Suomi F, Stenlund H, Johansson AI, Ehsan H, Salo VT, Montava-Garriga L, Naddafi S, et al. DGAT1 activity synchronises with mitophagy to protect cells from metabolic rewiring by iron depletion. EMBO J 2022; 41:e109390.

39. Nolan SJ, Romano JD, Coppens I. Host lipid droplets: An important source of lipids salvaged by the intracellular parasite Toxoplasma gondii. PLoS Pathog 2017; 13:e1006362.

40. Walther TC, Chung J, Farese RV. Lipid droplet biogenesis. Annu Rev Cell Dev Biol 2017; 33:491–510.

41. Krahmer N, Guo Y, Wilfling F, Hilger M, Lingrell S, Heger K, Newman HW, Schmidt-Supprian M, Vance DE, Mann M, et al. Phosphatidylcholine synthesis for lipid droplet expansion is mediated by localized activation of CTP:phosphocholine cytidylyltransferase. Cell Metab 2011; 14:504–15.

42. Romano JD, Sonda S, Bergbower E, Smith ME, Coppens I. Toxoplasma gondii salvages sphingolipids from the host Golgi through the rerouting of selected Rab vesicles to the parasitophorous vacuole. Mol Biol Cell 2013; 24:1974–95.

43. Pratt S, Wansadhipathi-Kannangara NK, Bruce CR, Mina JG, Shams-Eldin H, Casas J, Hanada K, Schwarz RT, Sonda S, Denny PW. Sphingolipid synthesis and scavenging in the intracellular apicomplexan parasite, Toxoplasma gondii. Molecular and Biochemical Parasitology 2013; 187:43–51.

44. Deevska GM, Nikolova-Karakashian MN. The expanding role of sphingolipids in lipid droplet biogenesis. Biochimica et Biophysica Acta (BBA) - Molecular and Cell Biology of Lipids 2017; 1862:1155–65.

45. Coppin A, Dzierszinski F, Legrand S, Mortuaire M, Ferguson D, Tomavo S. Developmentally regulated biosynthesis of carbohydrate and storage polysaccharide during differentiation and tissue cyst formation in Toxoplasma gondii. Biochimie 2003; 85:353–61.

46. Nolan SJ, Romano JD, Kline JT, Coppens I. Novel approaches to kill Toxoplasma gondii by exploiting the uncontrolled uptake of unsaturated fatty acids and vulnerability to lipid storage inhibition of the parasite. Antimicrob Agents Chemother 2018; 62.

47. Haschka D, Hoffmann A, Weiss G. Iron in immune cell function and host defense. Seminars in Cell & Developmental Biology 2021; 115:27–36.

48. Murdoch CC, Skaar EP. Nutritional immunity: the battle for nutrient metals at the host– pathogen interface. Nat Rev Microbiol 2022; 20:657–70.

49. Andreini C, Putignano V, Rosato A, Banci L. The human iron-proteome. Metallomics 2018; 10:1223–31.

50. Ying Z, Yin M, Zhu Z, Shang Z, Pei Y, Liu J, Liu Q. Iron stress affects the survival of Toxoplasma gondii [Internet]. Research Square; 2023 [cited 2023 Nov 30]. Available from: https://www.researchsquare.com/article/rs-3240882/v1

51. Sloan MA, Scott A, Harding CR. Keeping FIT: Iron-mediated post-transcriptional regulation in Toxoplasma gondii [Internet]. BioRxiv; 2023 [cited 2023 Nov 30]. Available from: http://biorxiv.org/lookup/doi/10.1101/2023.11.08.565792

52. Lüder CGK, Rahman T. Impact of the host on Toxoplasma stage differentiation. Microb Cell 2017; 4:203–11.

53. Walsh D, Katris NJ, Sheiner L, Botté CY. Toxoplasma metabolic flexibility in different growth conditions. Trends in Parasitology 2022; 38:775–90.

54. Blume M, Seeber F. Metabolic interactions between Toxoplasma gondii and its host. F1000Res 2018; 7.

55. Hu X, Binns D, Reese ML. The coccidian parasites Toxoplasma and Neospora dysregulate mammalian lipid droplet biogenesis. J Biol Chem 2017; 292:11009–20.

56. Gomes AF, Magalhães KG, Rodrigues RM, De Carvalho L, Molinaro R, Bozza PT, Barbosa HS. Toxoplasma gondii-skeletal muscle cells interaction increases lipid droplet biogenesis and positively modulates the production of IL-12, IFN-g and PGE2. Parasites Vectors 2014; 7:47.

57. Jarc E, Petan T. Lipid droplets and the management of cellular stress. Yale J Biol Med 2019; 92:435–52.

58. Konrad C, Wek RC, Sullivan WJ. GCN2-like eIF2α kinase manages the amino acid starvation response in Toxoplasma gondii. Int J Parasitol 2014; 44:139–46.

59. Augusto L, Martynowicz J, Amin PH, Carlson KR, Wek RC, Sullivan WJ. TgIF2K-B is an eIF2α kinase in Toxoplasma gondii that responds to oxidative stress and optimizes pathogenicity. mBio 2021; 12:e03160–20.

60. Augusto L, Wek RC, Sullivan WJ. Host sensing and signal transduction during Toxoplasma stage conversion. Mol Microbiol 2021; 115:839–48.

61. Bilgic B, Pfefferbaum A, Rohlfing T, Sullivan EV, Adalsteinsson E. MRI estimates of brain iron concentration in normal aging using quantitative susceptibility mapping. Neuroimage 2012; 59:2625–35.

