## Supplemental Figures S1 & S2 for "Iron depletion has different consequences on the growth and survival of *Toxoplasma gondii* strains"

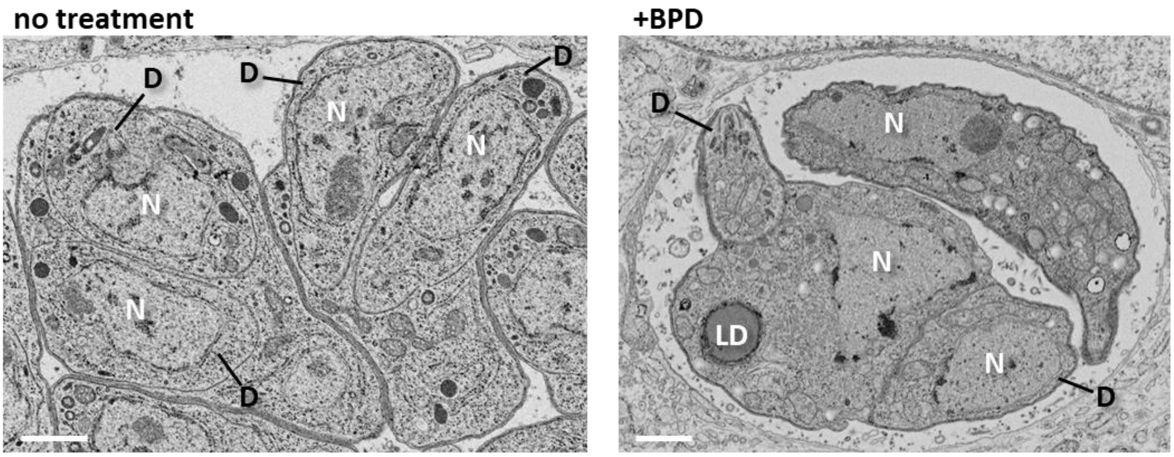


**Figure S1.** BPD treatment impairs daughter cell budding. The left electron microscopy image shows untreated RH parasites dividing almost synchronously, awhith each daughter cell (D) having incorporated organelles like the nucleus (N). The left electron microscopy picture shows RH parasites after a two days BPD treatment, displaying asynchronous division and daughter cell budding leaving out organellar material. LD: lipid droplet. Scale bar = 1 µm.

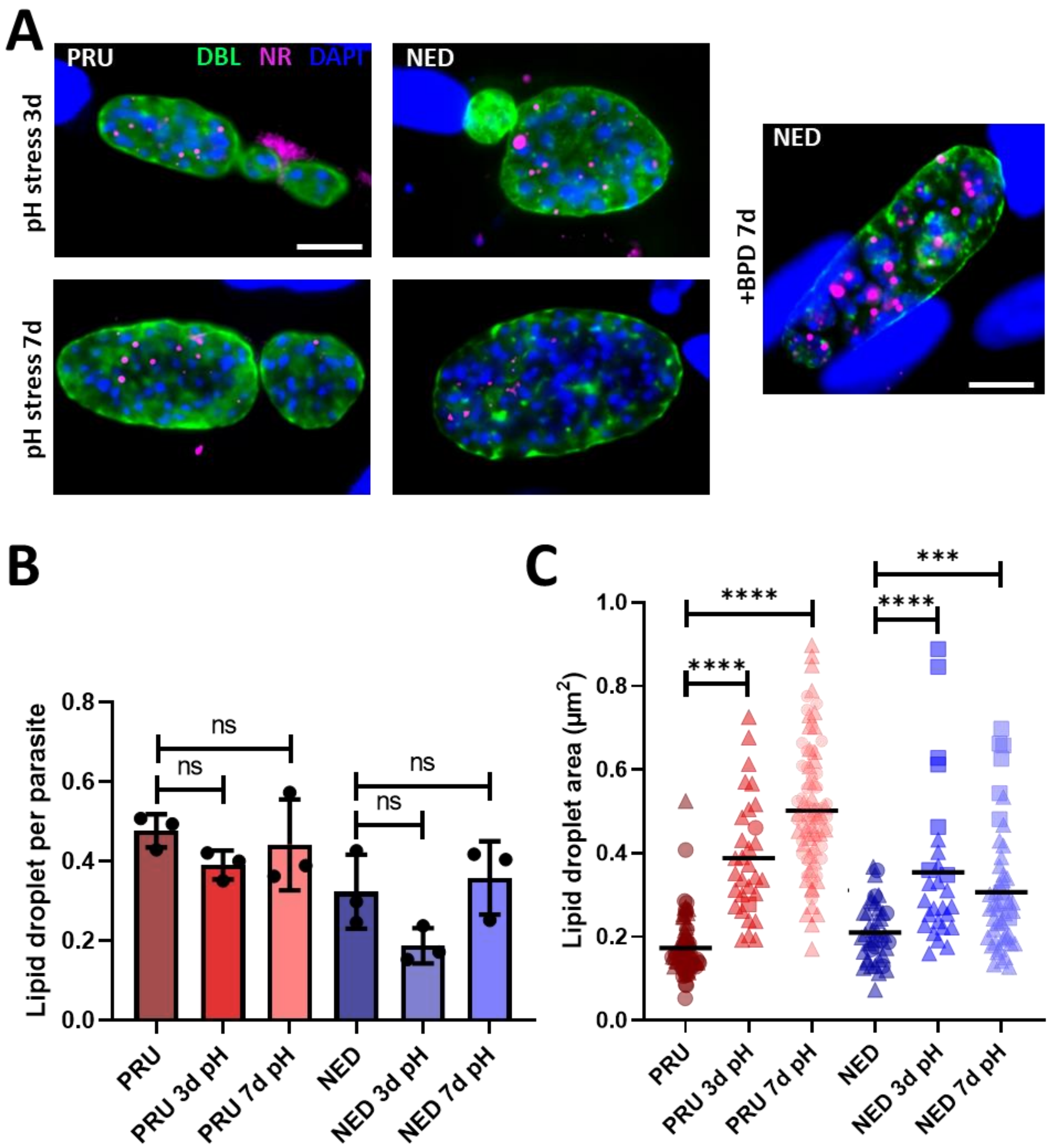


**Figure S2**. Lipid droplets in alkaline pH-induced cysts. **A.** Parasites of the type II and III cystogenic strains were submitted to alkaline pH-induced differentiation for 3 or 7 days and imaged for the cyst wall (DBL) and for lipid droplet (LD) content with Nile red (NR). Representative image of a cyst with type III parasites obtained after 7 days of BPD treatment is shown on the right for comparison. DNA was stained with DAPI. Scale bars = 10 µm. **B.** Quantification of LD numbers per parasite after inducing or not conversion into bradyzoites by pH stress for 3 or 7 days. Data are mean values ± SD from $n = 3$ independent experiments. At least 280 parasites were counted in each experimental condition. ns: not statistically significant, Student's *t*-test. **C.** Measurement of LD area in parasites after inducing or not conversion into bradyzoites by pH stress for 3 or 7 days. Data are mean values from $n = 3$ independent experiments. At least 40 LDs were measured in each experimental condition. Symbols are matched between identical experimental groups. *** $p \leq 0.001$, **** $p \leq 0.0001$, non-parametric Mann-Whitney test.
